# Novel use of Diffusion Tensor Imaging to Delineate the Rat Basolateral Amygdala

**DOI:** 10.1101/439836

**Authors:** Andre Obenaus, Eli Kinney-Lang, Amandine Jullienne, Elizabeth Haddad, Duke Shereen, Ana Solodkin, Jeffery F. Dunn, Tallie Z. Baram

**Affiliations:** Department of Pediatrics, Loma Linda University School of Medicine, Loma Linda, CA, 92350, USA; Departments of Anatomy/Neurobiology; University of California Irvine, Irvine, CA, 92697-4475, USA; Departments of Pediatrics; University of California Irvine, Irvine, CA, 92697-4475, USA; Departments of Neurology, University of California Irvine, Irvine, CA, 92697-4475, USA; Department of Radiology, Hotchkiss Brain Institute, Cumming School of Medicine, University of Calgary, Calgary, Alberta, T2N 4N1 Canada.

**Keywords:** Gadolinium, magnetic resonance imaging, rodent, volumes

## Abstract

The amygdaloid complex, including the basolateral nucleus (BLA) contributes crucially to emotional and cognitive brain functions, and is thus a major target of research in both humans and rodents. However, delineating structural amygdala plasticity in both normal and disease-related contexts using neuroimaging has been hampered by the difficulty of unequivocally identifying the boundaries of the BLA. This challenge is a result of poor contrast between BLA and the surrounding gray matter, including other amygdala nuclei. Here we describe a novel DTI approach to enhance contrast, enabling optimal identification of BLA in rodent brain from MR images. We employed this methodology together with a slice-shifting approach to measure BLA volume. We then validated the results by direct comparison to both histological and cellular-identity (parvalbumin)-based conventional techniques for defining BLA in the same brains used for MRI. We also confirmed the BLA region using DTI based tractography. The novel approach used here enables accurate and reliable delineation of BLA. Because this nucleus is involved in, and is changed by, developmental, degenerative and adaptive processes, the instruments provided here should be highly useful to a broad range of neuroimaging studies. Finally, the principles used here are readily applicable to numerous brain regions and across species.

**Summary Statement:** Use of MRI directionally encoded diffusion tensor imaging (DTI) can delineate the basolateral amygdala (BLA) and volumes derived from DTI were found to match those obtained using histological methods. Our approach can be used to identify the BLA.

## Introduction

The mammalian amygdala plays critical roles in emotional processing, fear, motivation and attention along with learning and memory. The amygdala itself is a constellation of nuclei and cell types that are remarkably conserved among mammalian species including rat, monkey and human (Chareyron et al., 2011). The amygdala is primarily composed of the basolateral (BLA) amygdala (lateral, basal, accessory basal nuclei), the central, medial and cortical nuclei, which have been termed the amygdaloid complex (Pitkanen et al., 1997; LeDoux, 2007). Among these nuclei, the BLA is thought to be integral to the processing of affective memories (Ritov et al., 2014). In humans, connectivity studies using functional magnetic resonance imaging (fMRI) and intracranial recording studies have demonstrated that the BLA complex is integral to selective memory formation and retrieval, and affective processing (Murray et al., 2014). Similar findings in the rodent have shown its importance in associative learning and fear processing (Roozendaal et al., 2002; Duvarci and Pare, 2014).

The amygdala figures prominently in several human disease states, including temporal lobe epilepsy and several types of dementia, as well as in major psychiatric disorders. Indeed, changes in amygdala volume have been reported in depression (Drevets, 2000; McEwen, 2001). The importance of the amygdala and particularly the BLA in normal brain function and in neuropsychiatric diseases demands establishment of imaging tools that can clearly delineate its anatomical boundaries.

Anatomical studies of the rodent amygdala have provided details about its cellular composition, connectivity and functional roles (Kemppainen and Pitkanen, 2000; Pitkanen et al., 2000; Shammah-Lagnado et al., 2001; Majak and Pitkanen, 2003; Unal et al., 2014). Human anatomical studies are few, and neuroimaging approaches have yielded a wealth of information. These neuroimaging studies range from volumetric assessments (Goncalves Pereira et al., 2005; O’Doherty et al., 2015; Lupton et al., 2016) to functional MRI studies (Manelis et al., 2015; Tajima-Pozo et al., 2016). In many of these studies both the volume and the location of the amygdala is based on atlas templates that provide general information, but do not provide details about the specific nuclei within the amygdala. Loss of detail is often the norm in imaging studies due to large slice thickness; however, the lack of gray matter contrast in the temporal lobe is the primary cause underlying the paucity of studies of the BLA. Recently, attempts have been made to resolve these issues using diffusion tensor imaging (DTI) approaches (Solano-Castiella et al., 2010; Saygin et al., 2011; Delgado y Palacios et al., 2014).

Similar to the human brain, traditional neuroimaging (T1, T2) of rodent amygdala has yielded poor differentiation between the amygdala and the surrounding structures, such as the piriform cortex and the caudate putamen. Even more problematic is differentiation of the BLA from other amygdala nuclei, such as the central and medial portion of the amygdala. Several strategies have been applied to enhance contrast and enable anatomy and functional connectivity, including manganese-enhanced imaging (MEMRI) (Aoki et al., 2004; Bangasser et al., 2013). Notably, these approaches have been largely unsuccessful in parcellating the specific amygdala nuclei, including the BLA. Others have utilized general delineation of the amygdala, again without distinction of its component nuclei (Bouilleret et al., 2009). More recently, DTI and its variants have been utilized to identify and visualize alterations, predominately in white matter in both rodents and humans (Wu and Cheung, 2010; Bracht et al., 2015). The use of DTI to probe grey matter microstructure are lacking, particularly in rodent studies. Finally, while rodent atlases are emerging with excellent definition of amygdala boundaries (Papp et al., 2014) many lack subdivisions of the amygdala such as BLA and related nuclei. Thus, there is a need for identifying reliable and robust methods for visualizing the individual nuclei of the amygdala and particularly the BLA.

We hypothesized that conventional imaging would not provide adequate delineation of the BLA but DTI and its ability to report microstructure could provide enhanced visual and quantitative assessments of the BLA. Thus, we report here on a novel strategy using high-resolution DTI combined with a slice-shifting approach to visualize and extract BLA volumes in the adult rat. The BLA boundaries identified using MRI-derived data were in excellent concordance with independent histological and immunohistochemical measures derived from the same brains. We further confirmed the accuracy of these BLA regions using DTI tractography. These findings delineate a strategy to accurately identify and quantify the BLA and can be applicable to additional brain regions as well as across species.

## Materials and Methods

All protocols were approved by the Loma Linda University Animal Health and Safety Committee and the University of California-Irvine IACUC, and are in compliance with federal regulations.

Adult Sprague Dawley rats (329.8 ± 3.3 g, Harlan; n=24) were allowed to recover in the vivarium for 5-7 days prior to perfusion fixation via transcardiac perfusion using 4% paraformaldehyde (PFA). Whereas imaging of either the brain in the cranial vault (Papp et al., 2014) or of the brain alone have been reported, brain-only samples have been used to generate atlases (Kovacevic et al., 2005). Therefore, we elected to remove the brains from the cranial vault. Brains were postfixed in 4% PFA, washed and stored at 4°C in 0.1M PBS/0.05% azide until imaging and histology. Together, these procedures reduced the potential for artifacts particularly at high field strengths. Prior to imaging, brains were placed in Fluorinert FC-770 to facilitate susceptibility matched imaging.

### T1-and T2-weighted Magnetic Resonance Imaging (MRI)

*Ex vivo* brains underwent high resolution T2-weighted anatomical MRI using an 11.7 T Bruker Avance scanner (Bruker Biospin, Billerica, MA, USA) The multiecho sequence had the following acquisition parameters: a 256×256 matrix, 25 slices covering the whole brain at 0.6 mm slice thickness, 2 cm field of view, repetition time/echo=2903/10.2 ms, 10 echos, two averages for a scan time of 25 min. This resulted in a 78×78×600 μm/pixel resolution.

A subset animals (n=6) underwent ultra-high resolution T1-weighted anatomical MRI using an 11.7 T Bruker Avance scanner (Bruker Biospin, Billerica, MA, USA). The sequence used was a 3D Rapid Acquisition with Relaxation Enhancement (3D RARE), with a 256^3^ matrix, 2 cm field of view and 78 μm slice thickness, repetition time/echo=2388/15 ms and a single average (total scan time=~5 hr). This resulted in a 78×78×78 μm/pixel isotropic resolution.

An additional subset of animals underwent T2 and T1 weighted imaging as described above but after post fixative incubation with Gd-DTPA (1.5%, Magnevist®) (n=3) or with Gd-DTPA (1.5%) mixed in with the PFA (n=3) and followed post fixation incubation for 3-5 days. Both groups were imaged using the same scan parameters. A control group without contrast (n=3) was also imaged in parallel.

### Diffusion tensor imaging (DTI)

High-resolution Diffusion Tensor Imaging (DTI) was performed on the same rats as reported for standard MRI (see above). DTI-MR images were acquired using a 9.4 T Bruker Biospin MR imaging system (Paravision 5.1). *Ex vivo* brains were carefully positioned inside a 5ml plastic syringe, and submerged in Fluorinert to eliminate any background noise and to increase signal to noise ratio. Each DTI acquisition consisted of 50 slices, 0.5 mm thick encompassing the entire brain, a 1.92×192 cm field of view with a data matrix size of 128×128 which was zero-filled to 256×256 matrix during image reconstruction. A four-shot echo-planar imaging (EPI) sequence was used to acquire four averages of diffusion weighted images with b values of 0 (5 images) and 3000 s/mm^2^ (30 images in non-collinear directions), a diffusion pulse width of 4 ms and interpulse duration of 20 ms with a repetition time of 12500 ms and an echo time of 36 ms for an acquisition time of ~2hr. DTI data were post-processed using DSI Studio (National Taiwan University: http://dsi-studio.labsolver.org). Raw Bruker data were imported into DSI Studio, where fractional anisotropy (FA), axial diffusivity (AD), radial diffusivity (RD) and trace (ADC) parametric maps were generated. In addition, primary, secondary, and tertiary diffusion eigenvalue (λ1, λ2, λ3) maps were calculated using FSL (http://fsl.fmrib.ox.ac.uk/fsl/).

For tractography the BLA was outlined on a single high CNR direction (direction 11, see Figure 3; Bregma: −3.14mm). Deterministic tractography was then performed using the following global parameters: angular threshold=60; step size=0.05mm; smoothing=0.60. The fiber threshold was optimized by DSI Studio to maximize the variance between the background and foreground and between subjects. One million seeds unrestricted seeds were placed for the entire brain and the tracts passing through the BLA were extracted.

Directionally specific DTI data were acquired at 11.7T to accurately determine amygdala volumes (see below). Contrast to noise (CNR) ratios were calculated using the following equation: CNR=(siBLA − siCTX)/stdevNOISE, where siBLA is the signal intensity in the BLA, siCTX) is the signal intensity of the cortex directly above the hippocampus, and stdevNOISE is the standard deviation of the noise in an ROI outside the brain tissue (see Figure 3B). CNR ROIs were inserted from a template and only the BLA ROI was modified as needed to encompass its anatomical boundaries.

To obtain enhance volumetric accuracy whilst preserving signal to noise (SNR) and CNR ratios we used a slice-shifting method, using only a single directionally encoded DTI acquisition (direction 11). We used our standard DTI coronal slice thickness of 600 μm but after the initial acquisition we then shifted the entire slice packet by 200 μm in the axial direction. This shift was then repeated two more times. The standard DTI imaging sequence (see above) typically yielded 2-3 600 μm thick slices through the BLA. With the slice-shifting method, we obtained 6-9 slices encompassing the BLA with an effective inter-slice distance of 200 μm. The slices from each data set (3 data sets) were combined into a single file where each slice was anatomically contiguous using a custom Matlab routine (see Figure 5). This approach provided improved volumetric resolution.

### BLA volumetric analysis from DTI

We have defined basolateral complex of the amygdala (BLA) as composed of the basal, lateral, and accessory basal nuclei of the amygdala, as previously reported (Roozendaal et al., 2002). Manual regions of interest (ROI) were carefully drawn on the right and left BLA regions on slices encompassing the amygdala (see Figure 1) on DTI parametric maps based on known locations from anatomical atlases (Paxinos and Watson, 2006). The amygdala complex starts at approximately Bregma −1.30 and extends to −4.80 mm. The basal and lateral nuclei (BLA) are found within this anterior-posterior range from −1.40 to −3.80 mm. For our analysis the BLA was bounded by the external capsule a white matter tract that is readily discernible on MRI. The dorsal extent of the BLA starts at the level of the rhinal fissure and extends 1.5 to 2 mm ventrally. The BLA is also bounded by the striatal, cortical (piriform) and ventricular structures (central and medial nuclei of the amygdala) in the medial aspect. Volumetric analysis of total brain and BLA were performed on MR coronal slices using Cheshire image processing software (Hayden Image/Processing Group, Waltham, MA, USA). For the volumetric data, bilateral BLA boundaries were manually drawn on each data set from the slice-shifted directionally encoded DTI image series using the boundaries as described above. Areas from each slice were extracted and multiplied by the effective interslice distance (200 μM) and slice number to obtain BLA volumes. All data were extracted and summarized in Excel.

**Figure 1.**
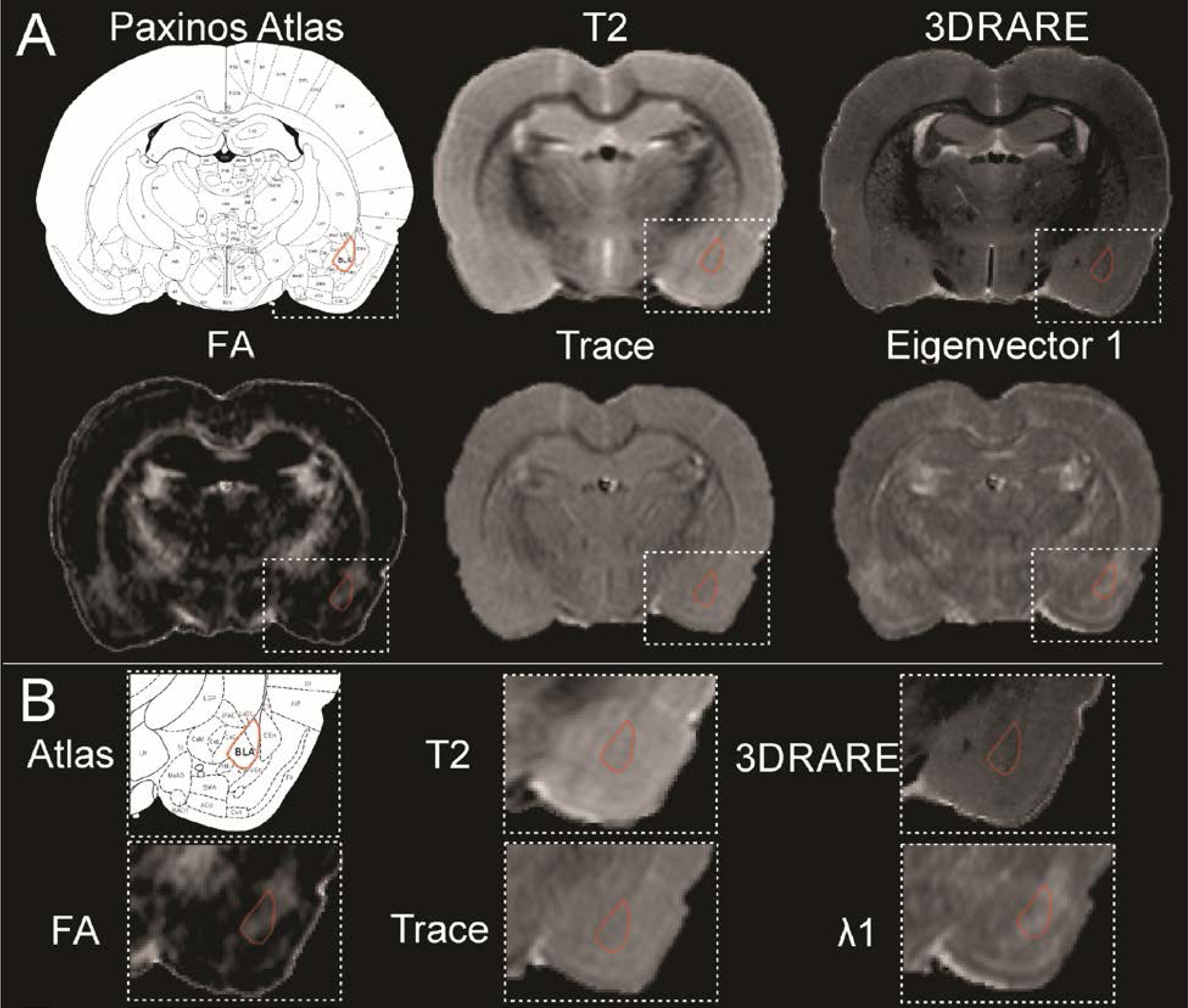
High resolution magnetic resonance imaging of the rodent brain does not elucidate the boundaries for the basolateral amygdala. A) The basolateral complex of the amygdala is highlighted (red) in several common imaging modalities including T2 weighted imaging, 3DRARE (3D Rapid Acquisition with Relaxation Enhancement) and diffusion tensor imaging parametric maps, including Fractional Anisotropy (FA), Trace and eigenvalue λ1. The anatomical boundaries of the BLA on the MR images were created by referencing the Paxinos Atlas. B) Expanded images from the MR images in A, further illustrates the difficulty in identifying the anatomical boundaries of the BLA. All images were collected from the same animal at 11.7T.

### Histology and Immunohistochemistry

After imaging was completed, brains were cryoprotected in 30% sucrose solution for 12 hours and frozen on dry ice. Coronal sections (30 μm) were cut and collected on Superfrost Plus (Fisher Scientific, Pittsburg, PA, USA) microscope slides (4 sections) and as free-floating sections stored in cryoprotectant (4 sections). This sequence was repeated every 120 μm. Tissue sections placed on slides were then processed for Cresyl violet (0.1%) staining as previously described (Obenaus et al., 2007).

Free-floating sections were processed for immunohistochemistry. The parvalbumin staining protocol was modified from Roozendaal et al (Roozendaal et al., 2002). Briefly, sections were removed from cryoprotectant solution and rinsed by 3 washes in PBS. Sections were then treated for 5 min with 3% H_2_O_2_ followed by non-specific sites blocking using 10% goat serum and 0.2% Triton X for 1.5h. Sections were incubated overnight with a parvalbumin antibody (1:200, #NB120-11427, Novus Biologicals, Littleton, CO, USA) at 4°C in the blocking buffer. After 3 rinses in PBS, sections were incubated with biotinylated goat anti-rabbit IgG antibody (1:500, #BA-1000, Vector laboratories, Burlingame, CA, USA) in PBS with 5% goat serum and 0.01% Triton X for 1h at room temperature. Following another 3 washes in PBS, sections were incubated in the avidin-biotin-peroxidase complex solution (Vector laboratories) for 30 min. Sections were then incubated in 0.012% 3,3’ diaminobenzidine solution containing 0.01% H_2_O_2_ to produce a brown reaction product. Sections were mounted on slides and dehydrated prior to being cover-slipped using Permount mounting medium (Fisher Scientific). For the MBP (myelin basic protein) staining, free-floating sections were washed, incubated in blocking buffer (2% goat serum in PBS) for 1.5 hours, then incubated overnight with anti-rat MBP antibody (1:250, MAB386, Millipore, Billerica, MA, USA) in PBS with 0.5% BSA. Alexa fluor 488 goat anti-rat IgG secondary antibody (1:1000, A-11006, Life Technologies, Carlsbad, CA, USA) was used for fluorescent detection of MBP.

Histological and immunohistochemical stained slides were imaged using a Keyence BZ-X700 microscope (Osaka, Japan) capturing the entire section at magnification 10X which was reconstructed using the XY-stitching feature. Total brain and BLA areas were manually delimited using anatomically defined boundaries (parvalbumin staining, internal capsule laterally, central nucleus medially and extending up ventrally to ~1.2mm from the base of the cortical tissues) and data was extracted from each section using BZ-X analyzer software (version 1.2.0.1) using the exact same ROI guidelines as those used for the MRI data. Histological BLA data were obtained by an investigator who did not undertake the MRI analysis.

### Statistical Analysis

All measurements and analyses were performed without knowledge of group. One-way analysis of variance (ANOVA), and Student’s *t*-tests were performed using GraphPad Prism5.0 (GraphPad, San Diego), followed by Bonferroni’s post-hoc tests when appropriate. Correlation analyses utilized were performed using matching data and were corrected using a Geisser-Greenhouse correction. Data are presented as mean ± SEM.

## Results

### Neuroimaging and contrast enhancement of the rodent amygdala

Given that the adult rat brain volume is ~2cc, we enhanced resolution through the use of high-field MRI. We eliminated motion artifacts and increased both contrast- and signal-to-noise ratios (CNR and SNR) by using *ex vivo* imaging. Both of these approaches aimed to improve visualization of the amygdala and its component nuclei. As noted in Figure 1 (A, B), conventional imaging (T2WI) resulted in ambiguous boundaries of the amygdala and in particular, of the BLA. For example, standard 2D T2-weighted imaging (78 μm in plane with 600 μm thick slices; n=3) provided reasonable visual contrast of many brain structures including white matter tracts and the hippocampal formation; however, contrast in the temporal regions including the amygdaloid complex remained poor.

### Exogenous contrast does not enhance definition of the BLA

Others have enhanced contrast among brain structures using exogenous contrast agents including gadolinium-based compounds. Accordingly, we employed two distinct approaches using Gd-DTPA, perfusion and incubation, in an attempt to increase contrast in the amygdala and visualize its nuclei (Figure 2). Whereas there was significant enhancement of signal to noise ratios within the amygdala and the temporal lobe (PFA:35.0, PFA, Gd incubation: 43.7, PFA+Gd, Gd incubation: 40.7) this strategy did not directly improve visualization of amygdala nuclei (data not shown). The method did enhance contrast in hippocampus and its sub-layers, as reported by others (Aoki et al., 2004; Bangasser et al., 2013).

**Figure 2.**
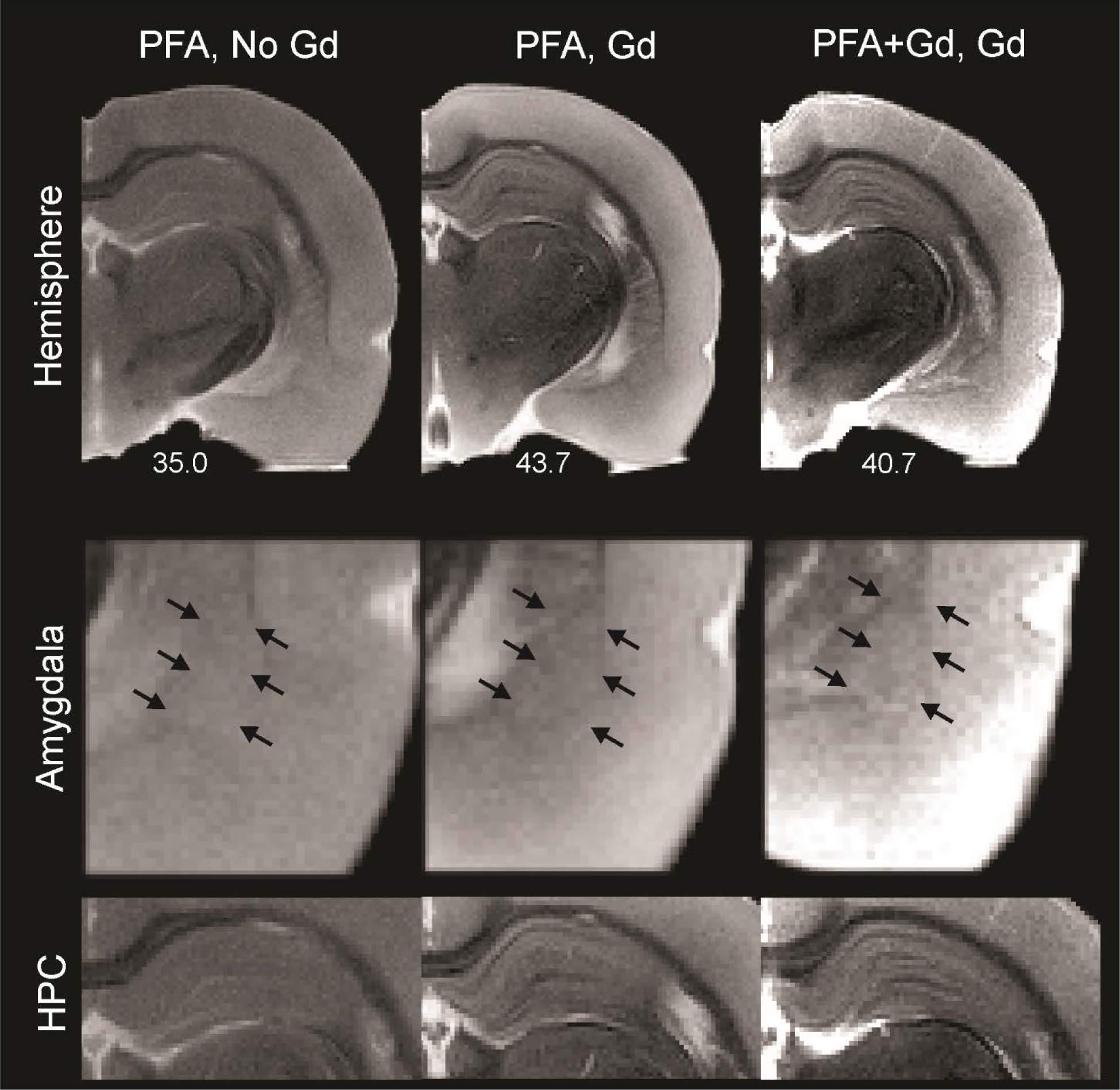
Exogenous contrast does not improve amygdala visualization. Exogenous contrast enhancement with Gadolinium (Gd), either within the perfusate or by immersion of *ex vivo* brain tissues did not dramatically improve visualization of the amygdala or the BLA (middle panel, arrows indicate the approximate location of the BLA). In contrast and similar to other published reports the hippocampus (HPC) exhibited enhanced contrast under both treatment paradigms (bottom panel). The signal-to-noise ratio values were improved with Gd treatment (values are reported under the hemispheres).

**Figure 3.**
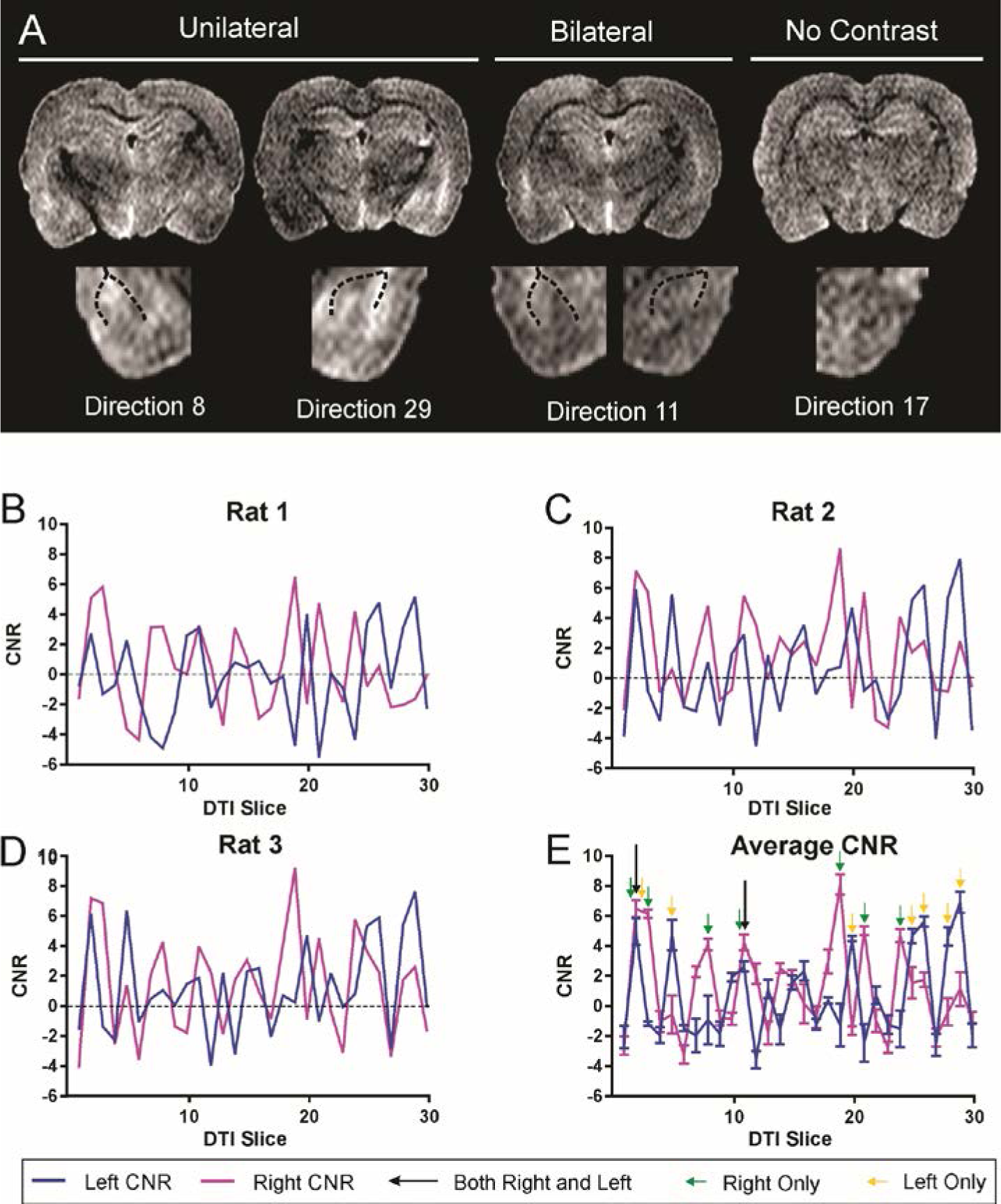
Optimal diffusion tensor directionality for identification of the basolateral complex of the amygdala. A) Examples of directional diffusion tensor images wherein either left, right (unilateral) or bilateral BLA appears to have increased contrast. A tensor image in which there is no enhanced contrast within the BLA is also shown. The dotted lines demarcate the external white matter boundaries of the external capsule that bound the BLA in the temporal lobe. See also Supplemental Figure 1. B) Contrast to noise ratio (CNR) measures at the level of the BLA (Bregma, −3.30) from each directional vector (see A) demonstrates remarkable consistency between three animals (A-C) in enhanced contrast in directional tensor images. CNR measures were then averaged across all three brains to identify the optimal unilateral or biolateral CNR in the BLA. Green arrows indicate optimal CNR in the right BLA while yellow arrows identify the optimal directions for the left BLA. Two directions (2, 11) have optimal CNR in both BLA simultaneously. While there are optimal directions for either the left or right BLA only, there are two directional DTI images which produces the highest CNR for both the left and right basolateral amygdala within the same MR image and with the least variance (direction 2 and 11). See also Supplemental Figure 2.

### High resolution neuroimaging and conventional DTI do not enhance BLA definition

Since standard neuroimaging, either with or without contrast, did not appreciably improve discrimination of rodent amygdala and its nuclei from the surrounding gray matter, we opted to amplify MR resolution. However, high resolution isotropic volumetric acquisitions (3DRARE; 78 μm isotropic, 5 hr acquisition) did not provide any additional visual improvements in contrast of the BLA (Figure 1A, B). Therefore, we turned to DTI as an alternative method for enhancing contrast (Wu and Cheung, 2010). We acquired DTI data using a scheme encompassing 30 directional vectors (78 μm in plane, 600 μm thick slices) (n=3). However, the anatomical boundaries of the amygdala were not appreciably enhanced visually nor better delineated on the DTI derived parametric maps, including the resultant FA, trace or eigenvalue images, in comparison to routine T1 or T2 imaging (Figure 1 A, B).

### Specific directionally-encoded DTI directions enhance BLA visualization

Whereas the DTI aggregate parametric maps did not improve visualization of the BLA, we reasoned that specific directional encoding vectors might yield enhanced CNR between BLA and surrounding structures. Such an approach has not been previously reported. To test this hypothesis, we used known anatomical landmarks and the enhanced CNR of the DTI images to draw the BLA ROIs on a single slice (Bregma=−3.30 mm) on each of the directionally encoded DTI image and the CNR was calculated (Figure 3). An example of all the 30 directional DTI images is shown in Supplemental Figure 1.

Visual examination of directionally encoded images at the level of the BLA revealed that the structure could be more readily observed in some directions compared to others (Figure 3A). Indeed, in some images the enhanced visualization were only unilateral and in select directions the BLA could be easily observed bilaterally. In contrast, many directions had poor or no observable BLA boundaries.

Analyzing CNR independently for each of the individual directional DTI vectors, we identified several vectors that yielded CNR signals that were greater than 4. Specifically, directions 2 and 11 had CNR of 4.93 and 6.47, respectively which provided optimal contrast simultaneously in both right and left BLA (Figure 3B, Supplemental Figure 2). Interestingly, we also observed vectors that provided increased CNR (4.08 to 8.10) for either the right or the left BLA, (right-direction 2, 3, 8, 11, 19, 21, 24; left direction 2, 5, 20, 25, 26, 28, 29). This finding of unequal contrast between the right and left BLA is expected, because of the different intrinsic properties of the brain structures lying either medial or lateral to the BLA. These features are the source of the contrast uniquely on the left or on the right amygdala for each individual directional vector. In addition, due to the in-homogenous cellular composition of the BLA itself, diffusion differences may exist along its left and right borders. Considering each side separately, CNR in the side-specific directions ranged from 4.08 to 8.10, a range that enhanced visualization. However, only directional vector images that contained simultaneous increased CNR in both BLA regions were utilized for volumetric analysis.

To further confirm our CNR approach for identifying optimal DTI vectors we undertook additional CNR measures of the adjacent striatal region. We observed increased CNR (2.40 to 5.01) in directions similar to those seen in the cortical CNR results. The increased CNR exhibited 80% concordance in the right hemisphere (right-direction 3, 8, 19, 21, 24) and 83% in the left hemisphere (left direction 5, 20, 22, 26, 28, 29). The decreased CNR values from the striatum are expected due to its increased iron content.

We then tested if the approach described above could facilitate volumetric analysis of the BLA. Plasticity and volume changes of BLA, for example as a result of augmented dendritic branching, have been described using postmortem methodologies (McEwen and Chattarji, 2004). However, assessment of BLA volume using imaging methods have been hampered by MRI slice thicknesses (often >500 μm) required for sufficient SNR within the context of a reasonable DTI acquisition timeframe. Notably, adult rat amygdala is only ~1000 μm anterior-posterior, so that a slice thickness of 500 μm would theoretically yield only two slices and thus resulting in insufficient resolution for accurate volumetry.

### Use of DTI and slice-shifting for accurate measures of BLA volume

We combined DTI, and specifically acquired data utilizing the directional encoding vector 11 (Figure 4; n=9) and sections that were acquired at 600 μm thickness. We then shifted the acquisition by 200 μm, then repeated this shift twice more. This resulted in an effective slice thickness of 200 μm (Figure 4A, Supplemental Figure 3). This approach yielded 6-9 slices (Figure 4A, right panel) that contained the BLA, in contrast to the 2-3 slices (Figure 4A, left panel) obtained using routine 600 μm thick section acquisitions. From these slice shifted data we calculated a BLA volume of 1.44 ± 0.02 and 1.47 ± 0.04 mm^3^ (mean ± SEM for the right and left BLA, respectively (Figures 4, 5, 6). The MRI-based volumes of right and left BLA volumes were not significantly different from each other (p=0.26) (Figure 4C). Combined BLA volume (right and left combined) was 2.94 ± 0.05 mm^3^.

**Figure 4.**
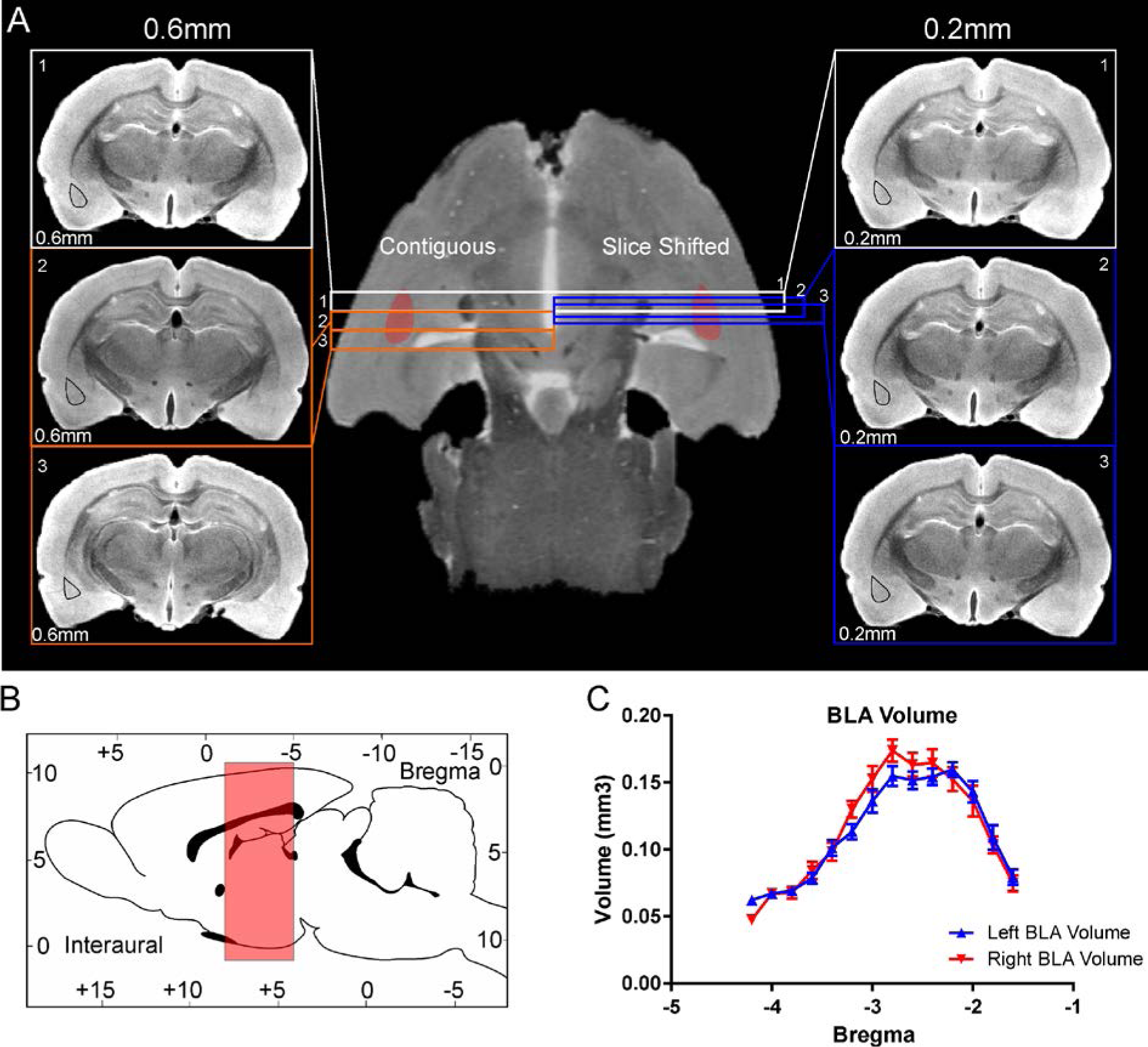
Basolateral amygdala volumes derived using a slice-shifting approach. A) Conventional sequential DTI of the rat brain utilized a slice thickness (contiguous) of 0.6 mm for optimal signal to noise/contrast to noise (SNR/CNR). Thus, with this acquisition method small anatomical structures, such as the BLA, only 2-3 slices capture the volume of the BLA (left-hand panel, slice thickness 0.6 mm, isodistance between slices = 0.6 mm, left BLA region outlined in black). In contrast, a slice-shifting technique that retained the same slice thickness (for optimal SNR/CNR) but captured a series of slices through the BLA at different isodistances with offsets of 0.2 mm (right-hand panel, slice thickness 0.6 mm, isodistance between slices = 0.2 mm) provided 6-9 slices encompassing the BLA. Using slice-shifting, the small area of the basolateral complex of the amygdala is captured in a greater number of slices for accurate volumetric analysis. B) Sagittal schematic of the rat brain illustrating the small region encompassed by the amygdala (red rectangle) from which DTI data were obtained. C) The slice shifting method yields an average BLA volume in an anterior-posterior direction illustrates similar right and left BLA volumes that concur with histology and volumes reported in the literature (see Figure 5). See also Supplemental Figure 3.

**Figure 5.**
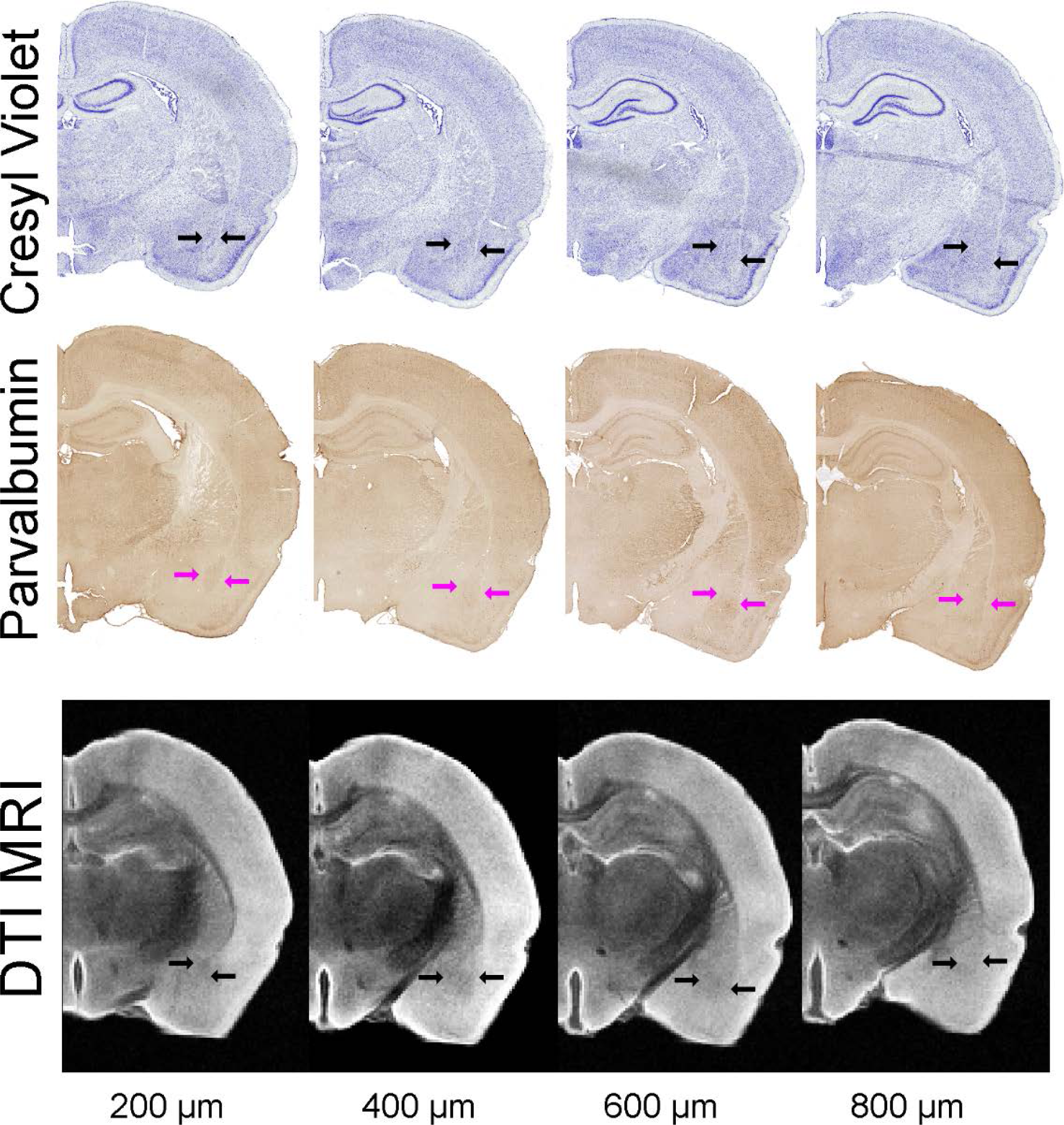
Histological delineation of the BLA. Histological sections (30 μm thick) were matched to MRI DTI slices (direction 11, 200 μm thick) for volumetric analysis. Cresyl Violet staining and parvalbumin immunohistochemistry for BLA were utilized to derive BLA volumes. Blue and magenta arrows outline the BLA on the histology. The DTI eigenvalue images are from direction 11 from slices 200 μm apart. All data in this figure are derived from the same animal.

### Histological comparison and validation of MRI-derived BLA volumes

To examine the accuracy and utility of the MRI-derived delineation and volume measurements of the BLA, we undertook direct comparison of the DTI method to two independent measurements performed directly on the same brain used for imaging. Representative histological samples from the same animal are shown in Figure 5. Cresyl violet staining of tissue sections encompassing the BLA resulted in volumes for right and left BLA of 1.36 ± 0.04 and 1.40 ± 0.06 mm^3^, respectively, and a combined volume of 2.76 ± 0.06 mm^3^ (Figures 5, 6A-C). The BLA contains large numbers of parvalbumin (PV) containing cells, and the presence of these neurons has been used to delineate the boundaries of this nucleus (Roozendaal et al., 2002). Employing PV immunohistochemistry resulted in a calculated BLA volume for right and left (1.45 ± 0.07, 1.41 ± 0.10 mm^3^) and combined volume of 2.87 ± 0.14 mm^3^ (Figures 5, 6A, B, D).

**Figure 6.**
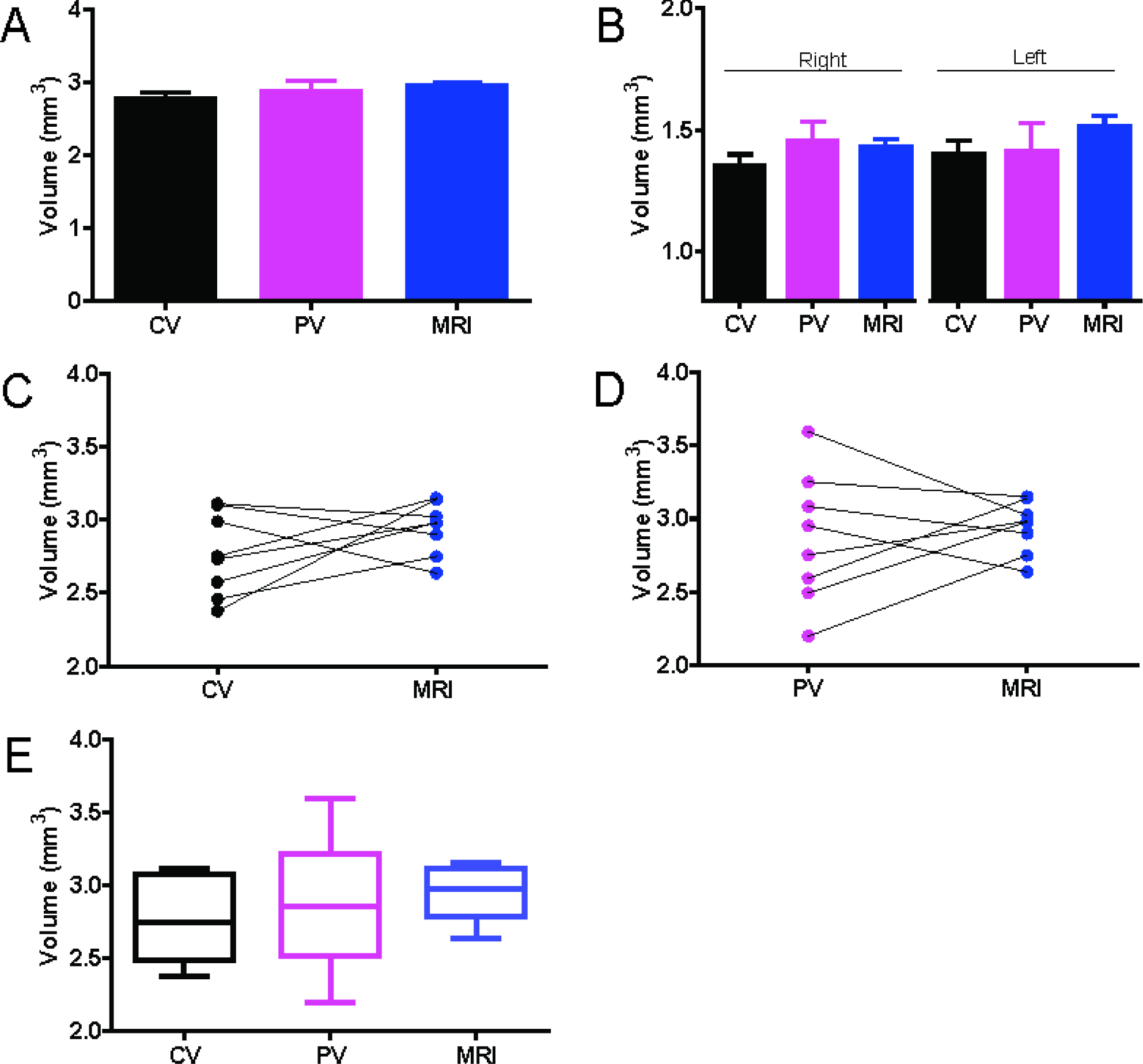
Histology and MRI BLA volumes. A) No significant differences were found in BLA volumes extracted from Cresyl violet (CV), parvalbumin (PV) or DTI MRI. B) Similarly, no significant differences were seen in volumes from either the right or left BLA. C) CV anatomical data matched well to MRI. D) PV data exhibited greater variability than CV in BLA volumes but matched MRI derived data. E) Box and whisker plots illustrate that BLA volumes from MRI had the lowest variability while PV exhibited the largest. This low variability in the MRI may be due to use of the entire data (6-9 slices), while histology extracted area from sections at 120 μm intervals.

We then examined statistically the results of the three methods of measuring BLA volume, focusing on comparison of the MRI-derived technique to the two direct tissue approaches. In essence, all methods yielded the same values. Specifically, Cresyl violet and PV volumes were comparable (p=0.60) for each side separately and for the combined BLA volume (p=0.060) (Figure 6C-E). BLA volumes derived from the two histological methods did not differ appreciably from those derived from DTI / slice-shifting (p=0.06, ANOVA). This was true also when each side was analyzed separately (right p=0.20; left p=0.19) (Figure 6C, D). The volumes obtained using the three methods were not appreciably different not only when the means of volumes derived from each method were compared, but also when the comparison was among BLA volumes of an individual brain that were examined using MRI or each of the two histological methods.

Interestingly, the variance among BLA volumes in the group of animals was smallest using MRI-derived analyses (Figure 6E), whereas PV delineation yielded the highest variance. The PV variance could be due in part to the rigors of the immunohistochemistry staining process.

To further demonstrate the robustness of the described approach, we undertook interrater calculations of the amygdala volumes from a random sub-set of three animals (Figure 6F-I). Rater A derived the original delineations reported herein, Rater B has extensive knowledge of MRI and the anatomy of the amygdala, and Rater C has limited experience with MRI or amygdala anatomy. Raters B and C were provided the manuscript and a coded dataset without additional training. As can be seen in Figure 6F-I, the experienced Rater (B) had volumes that were close to the originals whereas the Rater with the least experience (C) had the least similar volumes to the original. Further, MRI/amygdala experience appeared to reduced variance between animals (Figure 6G) with standard deviations of A: 0.047, B: 0.015, C: 0.528. Typically, Rater C overestimated the area on each slice of the MRI (Figure 6I). We then performed a training session for Rater C wherein volumes were now consistent with Raters A and B with a modest reduction in variability (A: 0.047, B: 0.015, C: 0.447). Volumes were not significantly different between raters A and C, but B was significantly smaller than A (p<0.0001). After training Rater C was not significantly different from Rater B (B:C1 p=0.046; B:C2 p>0.05; Figure 6G).

### Additional validation of MRI-derived BLA regions

To further validate our DTI derived approach for the BLA we performed two additional confirmatory measures. Firstly, we undertook immunohistochemical staining to elucidate the white matter tracts that are known to bound the BLA (Figure 7). Specifically, we used myelin basic protein (MBP) immunohistochemistry to identify the external capsule (EC) and the amygdalar capsule (AC). The BLA region we identified on high CNR DTI directions (e.g. direction 11) clearly match those seen and delineated by the white matter on MBP stained sections. This clear distinction was not evident in DTI low CNR directions (Figure 7).

**Figure 7.**
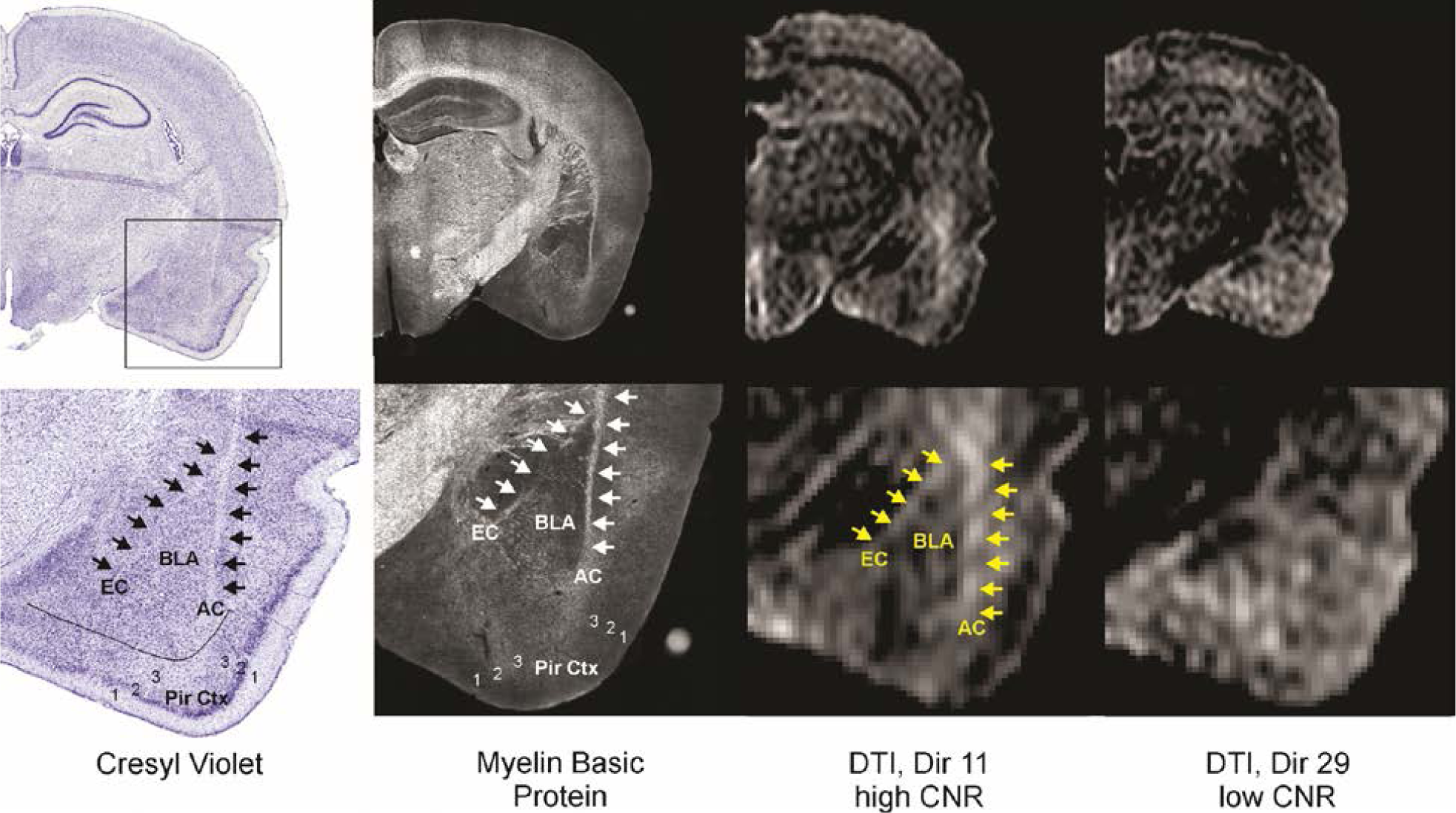
Further confirmation of the ability for DTI to delineate the BLA. To further demonstrate that specific directional DTI can delineate the BLA we compared cresyl violet and myelin basic protein (MBP) stained sections to high and low CNR DTI. The MBP staining was to provide further verification of known white matter tracts that bound the BLA, the external capsule (EC) and amgdalar capsule (AC). As can be appreciated, direction 11 which exhibited high CNR values exhibits clear boundaries of the BLA whereas a DTI direction with low CNR has little identifiable BLA landmarks. Arrows outline the predominate white matter tracts when observed. BLA - basolateral amygdala, EC - external capsule; white matter, AC - amygdalar capsule, white matter, Pir Ctx - piriform cortex where 1,2,3 identify the cortical layers. Histology is submicron/pixel resolution, whereas DTI is 75 μm/pixel in plane and 600 μm thick. (All images from the same animal).

Secondly, we undertook DTI tractography to validate the accuracy of our BLA boundaries. Whole brain tractography was performed followed by examining only those tracts that passed through the BLA region of interest base on our optimal CNR analysis (see Figure 3). The resultant tractography revealed a compact bundle of tracts that passed anteriorly from the olfactory bulbs through the BLA and on posteriorly to the ventral hippocampus with tracts looping anteriorly to the medial prefrontal cortex (mPFC). To further validate the accuracy of the BLA delineation we dilated the entire region of interest uniformly by 2 pixels which led to a dramatic increase in the number of unilateral tracts and the appearance of tracts crossing the midline (Figure 8B). A similar increase in tract density occurred when the original BLA regional designation was shifted leftward by 300 μm (4 pixels) (Figure 8C). The dramatic increase is due to incorporation of the amygdalar capsule white matter. A rightward shift of the BLA region by 300 μm also resulted in an increase in tracts, albeit less robust (Figure 8D) and encompasses the smaller white matter tract, the external capsule.

**Figure 8.**
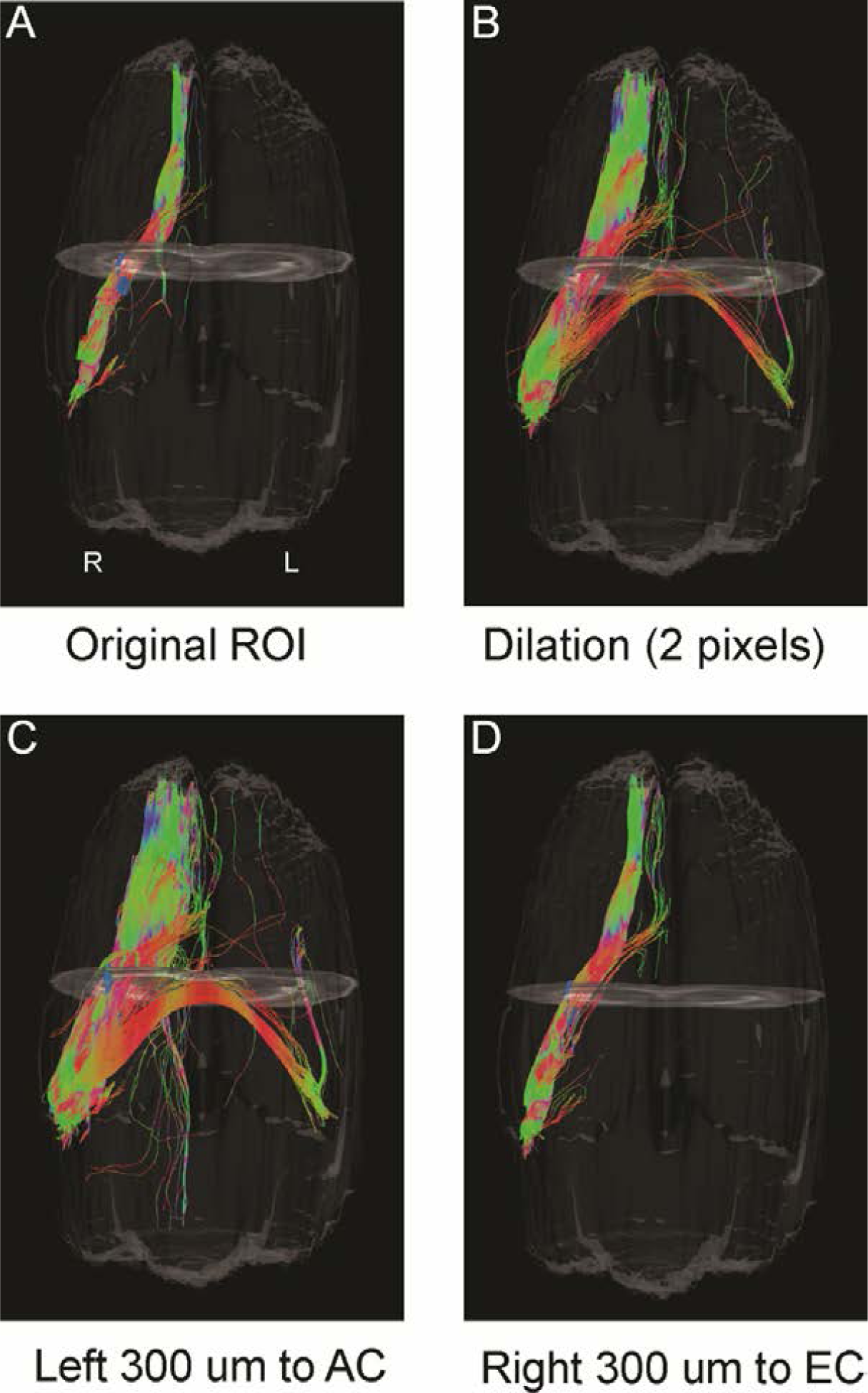
Tractography validation of the BLA regions from DTI. The BLA regions derived from high CNR directions were evaluated using tractography to evaluate known reciprocal connections between the medial prefrontal cortex (mPFC) and the BLA. A) Tracts (streamlines) from the using the BLA regions delineated on DTI direction 8 (high right CNR; see Figure 3E), illustrating a uniform and compact connection between the mPFC and the BLA. B) Uniform dilation by 2 pixels (150μm) of the BLA region resulted in a proliferation of tracts including bilateral projections. C) A leftward shift of BLA region by 300 μm dramatically increased the number of streamlines as well as the appearance of bilateral connections. This shift included portions of the amygdalar capsule. D) An identical right ward shift of 300 μm that now encompassed the smaller external capsule and potentially portions of the medial amygdala resulted in increased tract numbers.

## Discussion

The principal findings of these studies are: 1) high resolution DTI, employing specific, directionally-encoded vectors, enhances CNR and enables visualization of the BLA; 2) contrast to noise optimization for right and left amygdala can be accomplished independently, 3) coupled with slice-shifting, the method described here yields high concordance with histological BLA segmentation, allowing for accurate assessment of BLA volumes across time. Finally, the principles described here for DTI imaging of the amygdala in the rodent are applicable to other brain regions and across species.

We show here that high-field MR diffusion imaging permits resolution and delineation of BLA boundaries. This is important as volumetric increases or decreases of this structure have been reported in a broad range of rodent models that are used extensively to study the physiological importance of the amygdala in health and disease (Kemppainen and Pitkanen, 2000; Roozendaal et al., 2002; Majak and Pitkanen, 2003; Unal et al., 2014). Electrophysiology experiments as well as neuroanatomical studies can only be performed once, after the animal is dead (postmortem). There is a major need to provide an approach that allows for repeated and detailed evaluation of the BLA, yet, there have been no anatomically accurate approaches to assess volume of the BLA from MRI. MRI allows for repeated measures that are critical to assess volumetric changes with age, after a treatment, with disease progression or as a result of other manipulations.

To address this gap we show here that the use of specific directionally-encoded DTI results in maximal CNR that was used to enhance visualization of the BLA. Ours is the first to accurately report MRI derived volumes of the BLA with excellent concordance to histology. A number of previous studies have attempted to derive amygdala volumes from MRI (Bouilleret et al., 2009; Kassem et al., 2013; Spinelli et al., 2013) using standard imaging methods. These studies reported amygdala volumes ranging from 56-100 mm^3^ that are not consistent with those reported from anatomical studies (Chareyron et al., 2011). The principal issue is insufficient anatomical boundaries from which to assess amygdala volumes (Delgado y Palacios et al., 2014). Even, increasing resolution by increasing imaging times in *ex vivo* brains has not resulted in reliable amygdala volumes (Spinelli et al., 2013). DTI has been used previously to enhance visualization of the human (Entis et al., 2012; Solodkin et al., 2013) and rodent brain (Zalsman et al., 2015).

However, to our knowledge, DTI has not been used to directly estimate amygdala or BLA volumes in either species. Notably, our initial approach, using DTI parametric images did not provide enhanced visualization of the BLA nuclei.

Solano-Castiella and colleagues have reported that parametric maps derived from high field (3T) DTI could potentially subdivide the human amygdala into medial and lateral regions (Solano-Castiella et al., 2010; Solano-Castiella et al., 2011). Similar to our results, they noted poor contrast of the amygdala on FA images. However, color coded diffusional directions revealed sub-regions within the amygdala, but the authors noted that the relationship between these directional DTI clusters and their anatomical precision remains unexplored. These initial attempts in human studies, while important, emphasize the lack of BLA or other nuclei volumes from neuroimaging data. Interestingly, we did not observe enhanced amygdala structure in the rodent parametric maps (FA, radial, axial and mean diffusivity). However, by examining raw directionally encoded DTI images we found improved visualization in several of the DTI directional images. We used CNR to identify the best vector directions for BLA contrast bilaterally. We found several optimal directions based on CNR in both the right and the left BLA. Thus, features of the amygdala and its nuclei, along with their inherent anatomical connectivity and structure could be used to parcellate specific nuclei, such as BLA. Indeed, we demonstrated that DTI tractography of the BLA resulted in a compact set of tracts passing through the BLA regions we delineated and perturbation of the location of the BLA region resulted in dramatic increases in the number and distribution of tracts. Further, semi-automated or fully automated methods have been successfully used to segment brain regions primarily to enhance throughput of data analysis. We and others have used a variety of computational and computer vison approaches with success (Kruggel et al., 2008; Donovan et al., 2012; Ghosh et al., 2012; Bianchi et al., 2015) and similar approaches could be used to assist in automated segmentation of the BLA. A variety of clustering algorithms could also be utilized to delineate the BLA independent of anatomical landmarks.

We expanded our observation of increased CNR in the BLA on specific DTI directional vectors by using a slice-shifting technique to reliably capture the volume of the BLA whilst minimizing acquisition time but maximizing signal within the region of interest. Our MRI BLA volumes were found to be virtually identical on Cresyl violet staining and parvalbumin immunohistochemistry. Previous studies of BLA volumes from histology report a wide range of volumes from 0.35 to 2.39 mm^3^ (Goncalves et al., 2008; Pego et al., 2008; Chareyron et al., 2011). Our BLA volumes of 1.44-1.50 mm^3^ clearly fall within the range previously reported. Thus, we are confident that our novel DTI imaging strategy can be used to reliably report BLA volumes in health and disease across species.

There are a number of limitations of the current study. Firstly, the histological and MRI data were not corrected for tissue shrinkage after perfusion fixation but our BLA volumes are similar to other virtually identical histological studies (Chareyron et al., 2011). Further, since our MRI data were obtained from *ex vivo* brains after tissue fixation and we then performed cellular and immunohistochemical staining on the same brain tissue, altered tissue morphology is less of a concern. Secondly, we did not explicitly test our DTI slice-shifting approach *in vivo*, but we see no impediment to implementing this directly to *in vivo* rodents and to human data. Thirdly, in our current study we found two directionally encoded images that provided the optimum contrast for concurrent bilateral BLA visualization. It is likely that other DTI directions on other scanners with alternate encoding schemes will observe different enhancements compared to those reported herein. We do not perceive this as a significant obstacle for clinical translation nor basic science studies since examination of the optimal contrast is required only once for each encoding scheme from individual MRI scanners. However, it is important to note that our approach to BLA identification and volumes is sensitive to the encoding direction and the optimal directionally encoded DTI direction is likely to vary both by the number of encoding directions (i.e. 30 vs 60) and manner in which the gradients are applied by MRI hardware/software. Thus, the optimal CNR (see Figure 3) may have to be performed for each MR scanner (at least initially) or if the DTI acquisition scheme has been altered. A fourth limitation may be the effect of head motion, which could alter our proposed in vivo applicability. We have not explicitly tested the effects of motion or motion correction schemes in the current data, future work could evaluate the impact of motion on the sensitivity of our proposed method. Finally, in this study, manual regions of interest were performed because publically available MRI atlases do not have nuclear parcellation of the amygdala. Manual segmentation is not optimal for large scale studies and future work will aim to develop suitable atlases.

## Conclusions

The importance of the amygdala in emotional and affective signal processing has been well established. However, individual nuclei within the amygdala have specific processing roles and anatomical and volumetric delineation of these nuclei based on neuroimaging is becoming increasingly important. The poor MR contrast in the temporal lobe between the amygdala nuclei limits further non-invasive inquiry. While various strategies have been used to increase differentiation between these amygdala sub-structures, there has been limited success. To resolve this impasse in rodents, we utilized high resolution directionally encoded DTI, in combination with a slice-shifting approach. We delineated the BLA amygdala nucleus and calculated its volume. We confirmed our DTI derived volumes by comparing MRI results to two histological methods from the same animals and found excellent concordance. Thus, we have identified a novel approach using DTI directional encoded directions wherein there is increased contrast in the BLA (relative to the gray matter of the temporal lobe) combined with a slice-shifting approach to obtain reliable volume of the BLA. This approach is readily translatable to clinical and pre-clinical studies aiming to understand BLA role in in normal brain and in neuro-psychiatric disorders.

## Acknowledgements

Supported by NIH Grant P50MH096889 and NS35439 (TZB). We thank Mr. Tad Foniok, Mr. David Rushforth and Dr. Jeffery Dunn (University of Calgary) for assistance in acquiring the DTI data and funding from the Brain Canada Platform Grant.

The authors have no competing financial interests to declare.

## References

Aoki I, Wu YJ, Silva AC, Lynch RM, Koretsky AP (2004) In vivo detection of neuroarchitecture in the rodent brain using manganese-enhanced MRI. NeuroImage 22:1046–1059.

Bangasser DA, Lee CS, Cook PA, Gee JC, Bhatnagar S, Valentino RJ (2013) Manganese-enhanced magnetic resonance imaging (MEMRI) reveals brain circuitry involved in responding to an acute novel stress in rats with a history of repeated social stress. Physiology & behavior 122:228–236.

Bianchi A, Bhanu B, Obenaus A (2015) Dynamic low-level context for the detection of mild traumatic brain injury. IEEE transactions on bio-medical engineering 62:145–153.

Bouilleret V, Cardamone L, Liu YR, Fang K, Myers DE, O'Brien TJ (2009) Progressive brain changes on serial manganese-enhanced MRI following traumatic brain injury in the rat. Journal of neurotrauma 26:1999–2013.

Bracht T, Linden D, Keedwell P (2015) A review of white matter microstructure alterations of pathways of the reward circuit in depression. Journal of affective disorders 187:45–53.

Chareyron LJ, Banta Lavenex P, Amaral DG, Lavenex P (2011) Stereological analysis of the rat and monkey amygdala. The Journal of comparative neurology 519:3218–3239.

Delgado y Palacios R, Verhoye M, Henningsen K, Wiborg O, Van der Linden A (2014) Diffusion kurtosis imaging and high-resolution MRI demonstrate structural aberrations of caudate putamen and amygdala after chronic mild stress. PloS one 9:e95077.

Donovan V, Bianchi A, Hartman R, Bhanu B, Carson MJ, Obenaus A (2012) Computational analysis reveals increased blood deposition following repeated mild traumatic brain injury. NeuroImage Clinical 1:18–28.

Drevets WC (2000) Functional anatomical abnormalities in limbic and prefrontal cortical structures in major depression. Progress in brain research 126:413–431.

Duvarci S, Pare D (2014) Amygdala microcircuits controlling learned fear. Neuron 82:966–980.

Entis JJ, Doerga P, Barrett LF, Dickerson BC (2012) A reliable protocol for the manual segmentation of the human amygdala and its subregions using ultra-high resolution MRI. NeuroImage 60:1226–1235.

Ghosh N, Yuan X, Turenius CI, Tone B, Ambadipudi K, Snyder EY, Obenaus A, Ashwal S (2012) Automated core-penumbra quantification in neonatal ischemic brain injury. Journal of cerebral blood flow and metabolism : official journal of the International Society of Cerebral Blood Flow and Metabolism 32:2161–2170.

Goncalves L, Silva R, Pinto-Ribeiro F, Pego JM, Bessa JM, Pertovaara A, Sousa N, Almeida A (2008) Neuropathic pain is associated with depressive behaviour and induces neuroplasticity in the amygdala of the rat. Experimental neurology 213:48–56.

Goncalves Pereira PM, Insausti R, Artacho-Perula E, Salmenpera T, Kalviainen R, Pitkanen A (2005) MR volumetric analysis of the piriform cortex and cortical amygdala in drug-refractory temporal lobe epilepsy. AJNR American journal of neuroradiology 26:319–332.

Kassem MS, Lagopoulos J, Stait-Gardner T, Price WS, Chohan TW, Arnold JC, Hatton SN, Bennett MR (2013) Stress-induced grey matter loss determined by MRI is primarily due to loss of dendrites and their synapses. Molecular neurobiology 47:645–661.

Kemppainen S, Pitkanen A (2000) Distribution of parvalbumin, calretinin, and calbindin-D(28k) immunoreactivity in the rat amygdaloid complex and colocalization with gamma-aminobutyric acid. The Journal of comparative neurology 426:441–467.

Kovacevic N, Henderson JT, Chan E, Lifshitz N, Bishop J, Evans AC, Henkelman RM, Chen XJ (2005) A three-dimensional MRI atlas of the mouse brain with estimates of the average and variability. Cerebral cortex 15:639–645.

Kruggel F, Paul JS, Gertz HJ (2008) Texture-based segmentation of diffuse lesions of the brain’s white matter. NeuroImage 39:987–996.

LeDoux J (2007) The amygdala. Current biology : CB 17:R868–874.

Lupton MK et al. (2016) The effect of increased genetic risk for Alzheimer’s disease on hippocampal and amygdala volume. Neurobiology of aging 40:68–77.

Majak K, Pitkanen A (2003) Projections from the periamygdaloid cortex to the amygdaloid complex, the hippocampal formation, and the parahippocampal region: a PHA-L study in the rat. Hippocampus 13:922–942.

Manelis A, Ladouceur CD, Graur S, Monk K, Bonar LK, Hickey MB, Dwojak AC, Axelson D, Goldstein BI, Goldstein TR, Bebko G, Bertocci MA, Hafeman DM, Gill MK, Birmaher B, Phillips ML (2015) Altered amygdala-prefrontal response to facial emotion in offspring of parents with bipolar disorder. Brain : a journal of neurology 138:2777–2790.

McEwen BS (2001) Plasticity of the hippocampus: adaptation to chronic stress and allostatic load. Annals of the New York Academy of Sciences 933:265–277.

McEwen BS, Chattarji S (2004) Molecular mechanisms of neuroplasticity and pharmacological implications: the example of tianeptine. European neuropsychopharmacology : the journal of the European College of Neuropsychopharmacology 14 Suppl 5:S497–502.

Murray RJ, Brosch T, Sander D (2014) The functional profile of the human amygdala in affective processing: insights from intracranial recordings. Cortex; a journal devoted to the study of the nervous system and behavior 60:10–33.

O’Doherty DC, Chitty KM, Saddiqui S, Bennett MR, Lagopoulos J (2015) A systematic review and meta-analysis of magnetic resonance imaging measurement of structural volumes in posttraumatic stress disorder. Psychiatry research 232:1–33.

Obenaus A, Robbins M, Blanco G, Galloway NR, Snissarenko E, Gillard E, Lee S, Curras-Collazo M (2007) Multi-modal magnetic resonance imaging alterations in two rat models of mild neurotrauma. Journal of neurotrauma 24:1147–1160.

Papp EA, Leergaard TB, Calabrese E, Johnson GA, Bjaalie JG (2014) Waxholm Space atlas of the Sprague Dawley rat brain. NeuroImage 97:374–386.

Paxinos G, Watson C (2006) The Rat Brain in Stereotaxic Coordinates: Academic Press.

Pego JM, Morgado P, Pinto LG, Cerqueira JJ, Almeida OF, Sousa N (2008) Dissociation of the morphological correlates of stress-induced anxiety and fear. The European journal of neuroscience 27:1503–1516.

Pitkanen A, Savander V, LeDoux JE (1997) Organization of intra-amygdaloid circuitries in the rat: an emerging framework for understanding functions of the amygdala. Trends in neurosciences 20:517–523.

Pitkanen A, Pikkarainen M, Nurminen N, Ylinen A (2000) Reciprocal connections between the amygdala and the hippocampal formation, perirhinal cortex, and postrhinal cortex in rat. A review. Annals of the New York Academy of Sciences 911:369–391.

Ritov G, Ardi Z, Richter-Levin G (2014) Differential activation of amygdala, dorsal and ventral hippocampus following an exposure to a reminder of underwater trauma. Frontiers in behavioral neuroscience 8:18.

Roozendaal B, Brunson KL, Holloway BL, McGaugh JL, Baram TZ (2002) Involvement of stress-released corticotropin-releasing hormone in the basolateral amygdala in regulating memory consolidation. Proceedings of the National Academy of Sciences of the United States of America 99:13908–13913.

Saygin ZM, Osher DE, Augustinack J, Fischl B, Gabrieli JD (2011) Connectivity-based segmentation of human amygdala nuclei using probabilistic tractography. NeuroImage 56:1353–1361.

Shammah-Lagnado SJ, Alheid GF, Heimer L (2001) Striatal and central extended amygdala parts of the interstitial nucleus of the posterior limb of the anterior commissure: evidence from tract-tracing techniques in the rat. The Journal of comparative neurology 439:104–126.

Solano-Castiella E, Schafer A, Reimer E, Turke E, Proger T, Lohmann G, Trampel R, Turner R (2011) Parcellation of human amygdala in vivo using ultra high field structural MRI. NeuroImage 58:741–748.

Solano-Castiella E, Anwander A, Lohmann G, Weiss M, Docherty C, Geyer S, Reimer E, Friederici AD, Turner R (2010) Diffusion tensor imaging segments the human amygdala in vivo. NeuroImage 49:2958–2965.

Solodkin A, Chen EE, Van Hoesen GW, Heimer L, Shereen A, Kruggel F, Mastrianni J (2013) In vivo parahippocampal white matter pathology as a biomarker of disease progression to Alzheimer’s disease. The Journal of comparative neurology 521:4300–4317.

Spinelli S, Muller T, Friedel M, Sigrist H, Lesch KP, Henkelman M, Rudin M, Seifritz E, Pryce CR (2013) Effects of repeated adolescent stress and serotonin transporter gene partial knockout in mice on behaviors and brain structures relevant to major depression. Frontiers in behavioral neuroscience 7:215.

Tajima-Pozo K, Yus M, Ruiz-Manrique G, Lewczuk A, Arrazola J, Montanes-Rada F (2016) Amygdala Abnormalities in Adults With ADHD. Journal of attention disorders.

Unal G, Pare JF, Smith Y, Pare D (2014) Cortical inputs innervate calbindin-immunoreactive interneurons of the rat basolateral amygdaloid complex. The Journal of comparative neurology 522:1915–1928.

Wu EX, Cheung MM (2010) MR diffusion kurtosis imaging for neural tissue characterization. NMR in biomedicine 23:836–848.

Zalsman G, Gutman A, Shbiro L, Rosenan R, Mann JJ, Weller A (2015) Genetic vulnerability, timing of short-term stress and mood regulation: A rodent diffusion tensor imaging study. European neuropsychopharmacology : the journal of the European College of Neuropsychopharmacology 25:2075–2085.

